# Emergence of belief-like representations through reinforcement learning

**DOI:** 10.1101/2023.04.04.535512

**Authors:** Jay A. Hennig, Sandra A. Romero Pinto, Takahiro Yamaguchi, Scott W. Linderman, Naoshige Uchida, Samuel J. Gershman

**Affiliations:** Department of Psychology, Harvard University, Cambridge, MA, USA; Center for Brain Science, Harvard University, Cambridge, MA, USA; Department of Molecular and Cellular Biology, Harvard University, Cambridge, MA, USA; Future Vehicle Research Department, Toyota Research Institute North America, Toyota Motor North America Inc., Ann Arbor, MI, USA; Wu Tsai Neurosciences Institute, Stanford University, Stanford, CA, USA; Department of Statistics, Stanford University, Stanford, CA, USA

## Abstract

To behave adaptively, animals must learn to predict future reward, or value. To do this, animals are thought to learn reward predictions using reinforcement learning. However, in contrast to classical models, animals must learn to estimate value using only incomplete state information. Previous work suggests that animals estimate value in partially observable tasks by first forming “beliefs”—optimal Bayesian estimates of the hidden states in the task. Although this is one way to solve the problem of partial observability, it is not the only way, nor is it the most computationally scalable solution in complex, real-world environments. Here we show that a recurrent neural network (RNN) can learn to estimate value directly from observations, generating reward prediction errors that resemble those observed experimentally, without any explicit objective of estimating beliefs. We integrate statistical, functional, and dynamical systems perspectives on beliefs to show that the RNN’s learned representation encodes belief information, but only when the RNN’s capacity is sufficiently large. These results illustrate how animals can estimate value in tasks without explicitly estimating beliefs, yielding a representation useful for systems with limited capacity.

**Author Summary:** Natural environments are full of uncertainty. For example, just because my fridge had food in it yesterday does not mean it will have food today. Despite such uncertainty, animals can estimate which states and actions are the most valuable. Previous work suggests that animals estimate value using a brain area called the basal ganglia, using a process resembling a reinforcement learning algorithm called TD learning. However, traditional reinforcement learning algorithms cannot accurately estimate value in environments with state uncertainty (e.g., when my fridge’s contents are unknown). One way around this problem is if agents form “beliefs,” a probabilistic estimate of how likely each state is, given any observations so far. However, estimating beliefs is a demanding process that may not be possible for animals in more complex environments. Here we show that an artificial recurrent neural network (RNN) trained with TD learning can estimate value from observations, without explicitly estimating beliefs. The trained RNN’s error signals resembled the neural activity of dopamine neurons measured during the same task. Importantly, the RNN’s activity resembled beliefs, but only when the RNN had enough capacity. This work illustrates how animals could estimate value in uncertain environments without needing to first form beliefs, which may be useful in environments where computing the true beliefs is too costly.

## Introduction

One pervasive feature of animal behavior is the ability to predict future reward. For example, a dog may learn that when her owner picks up the leash, she is likely to be rewarded with a walk in the near future. In associative learning settings such as this one, animals learn to associate certain stimuli (e.g., her owner grabbing the leash) with future reward (e.g., a walk). The neural basis of associative learning has been interpreted through the lens of reinforcement learning (RL). In particular, one successful theoretical model posits that associative learning is driven by the activity of dopamine neurons in the midbrain, where spiking activity resembles the reward prediction error (RPE) signal in an RL algorithm called temporal difference (TD) learning [1, 2, 3, 4]. We will describe this algorithm in more detail below.

In many real-world scenarios, effectively predicting reward may require a deeper understanding of the structure of the world that goes beyond associating observations and reward. To continue the example above, suppose the dog’s owner keeps his car keys under the leash. Now if he picks up the leash, this does not necessarily mean he is about to take his dog on a walk. In other words, his intention to take his dog on a walk is now “partially observable.” Standard RL approaches are insufficient for learning in partially observable environments, as these methods assume that all relevant states of the environment are fully observable. One way to solve this problem is by using observations to form a Bayesian posterior estimate of each hidden state, called a *belief state* [5]. Future reward can then be estimated by applying standard RL methods like TD learning to belief states rather than the raw observations.

Do animals estimate future reward using belief states? Evidence for this idea is suggestive, although indirect. Previous experimental work has shown that the phasic activity of midbrain dopamine neurons resembles the RPEs of TD learning in partially observable environments, where TD learning is performed on belief states rather than observations [6, 7, 8, 9, 10, 11]. The brain may have dedicated machinery, perhaps in prefrontal cortex [12, 13], for computing belief states, which could then be provided to downstream areas, such as the basal ganglia, to perform standard RL algorithms such as TD learning [14]. This architectural division of labor resonates with the broader literature on probabilistic computation in cortex, which has identified several different ways in which belief states could be encoded by neural activity [15].

There are a few difficulties in using a belief state to solve RL tasks. First, the belief state assumes knowledge of the environment’s transition and observation dynamics—something that may be challenging for animals to acquire via observations alone. Indeed, there are well-documented examples of animals failing to learn or use the correct environment model [16]. Second, the belief representation does not scale well to more realistic tasks with higher-dimensional state spaces, as beliefs live in a continuous space whose dimensionality grows with the number of discrete states in the environment. Finally, the belief state includes knowledge about all states in the environment, regardless of whether those states are relevant to the task at hand. Luckily, one can often use approximate representations of beliefs to find solutions that work well in practice [17]. This suggests that there may be other representations, more compact than the full belief state, that are sufficient for the particular task of estimating future reward [18, 19].

To address these difficulties, here we take inspiration from deep reinforcement learning. In deep RL, rather than explicitly learning beliefs, an agent uses nonlinear function approximation to learn a hidden representation that is sufficient for performing the task [20]. Compared to the belief representation, this approach does not require explicit knowledge about the structure of the environment. It may also scale better to more complex tasks, by virtue of the agent not needing to represent any features of the environment that do not directly pertain to the task at hand. Because beliefs are a non-linear dynamical system, here we use recurrent neural networks (RNNs) as our nonlinear function approximator. This choice was also motivated by the observation from the machine learning literature that RNNs can perform well on complex partially observable tasks [21]. Previous work in computational neuroscience has explored whether RNNs can be used to directly compute beliefs [17]. Here, by contrast, we explore whether training RNNs in partially observable environments leads to their representations implicitly becoming more like beliefs.

We show that RNNs can be trained to perform two previously studied associative learning tasks [7, 10], reproducing experimentally observed dopamine neuron activity. We probe the representations learned by the RNNs, and show that the representations resemble beliefs from statistical, functional, and dynamical systems perspectives. Finally, we show how an RNN’s capacity determines the degree to which its representation resembles beliefs, without a concurrent impact on its ability to perform the task. These results illustrate how animals might estimate reward in partially observable environments without requiring any explicit representation of beliefs.

## Results

### Reinforcement learning in partially observable environments using belief states

The standard RL objective is to learn the expected discounted future return, or *value*, of each state:

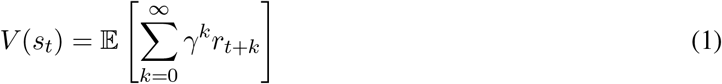

where *s*_*t*_ ∈ 𝒮 is the state of the environment at time *t*, 0 ≤ *γ <* 1 is a discount factor, and *r*_*t*_ is the reward. Rewards are random variables that depend on the environment state, and 𝔼 denotes an expectation over the potentially stochastic sequences of states and rewards. Because we are interested in modeling Pavlovian associative learning tasks, here we will assume there are no actions available to the agent. For notational simplicity, we will use the shorthand *V*_*t*_ = *V* (*s*_*t*_).

TD learning estimates the value function by exploiting a set of Markovian assumptions about the environment: the probability of state *s*_*t*_ depends only on the last state *s*_*t*−1_, and the probability of reward *r*_*t*_ depends only on the current state *s*_*t*_. Under these assumptions, the value function can be decomposed into a recursive form known as the *Bellman equation* [22]:

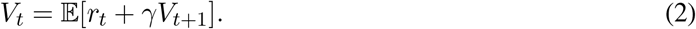

If an agent has access to an approximation of the value function, 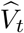, along with sample paths of states and rewards, then it can compute an estimate of the discrepancy between the approximate and true value function by taking the difference between the two sides of the Bellman equation:

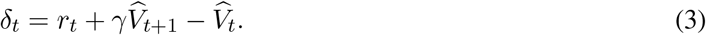

This is the temporal difference error, the precise definition of the RPE used in TD learning. Under the Bellman equation, 𝔼 [*δ*_*t*_] = 0 when 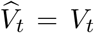. If 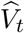 is determined by a set of adaptable parameters ***θ***, the approximation can be improved by following the stochastic gradient of the squared TD error:

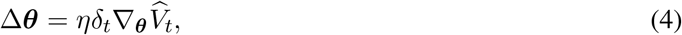

where 0 *< η <* 1 is a learning rate, and 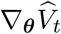 is the gradient of 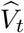 with respect to ***θ***. One common example is a linear function approximator:

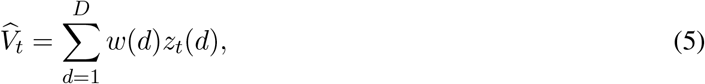

where *z*_*t*_(*d*) ∈ ℝ is some feature (indexed by *d*) of the state, and ***θ*** = ***w*** ∈ ℝ^*D*^ is a learned set of weights on all the features. Under this approximation, 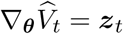.

In a partially observable Markov process, agents do not observe the state *s*_*t*_ directly, but instead observe only observations *o*_*t*_ ∈ 𝒪. Critically, observations are not in general Markovian, which means that TD learning methods cannot be naively applied to value function approximators defined over observations. One way of understanding this is to note that the value of an observation may depend on the long-term past [23]. In the dog leash example from the Introduction, the value (to the dog) of her owner picking up the leash depends on the history of events leading up to that moment—for example, if her owner recently announced that his car keys were missing. Reward prediction requires a compression of this history into a “sufficient statistic”—in effect creating a transformed state space over which the Markov property holds. TD learning can then be applied to this transformed state space.

One such transformed state is the posterior probability distribution over hidden states given the history of observations, also known as the *belief state* [5]:

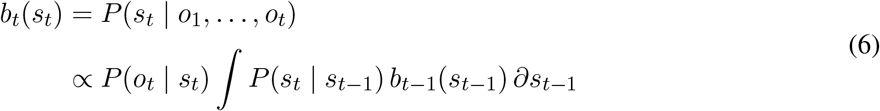

where the recursion follows from the Chapman-Kolmogorov equation, which stipulates how to update the belief state from *t* − 1 to *t* after observing observation *o*_*t*_.

For a partially observable Markov process with a finite state space, 𝒮= {1, …, *K*}, the value function can be written as a weighted combination of beliefs:

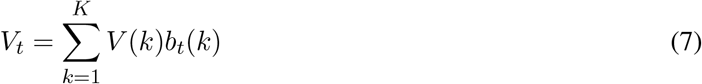

where *V* (*k*) and *b*_*t*_(*k*) are the value and belief of state *s*_*t*_ = *k*, respectively. This means that linear value function approximation in Eqn. (5) is sufficiently expressive for the partially observable problems we consider in this paper: with enough training data, the value function approximator will perfectly estimate the true value function (i.e., given learned weights *w*(*k*) = *V* (*k*) and features *z*_*t*_(*k*) = *b*_*t*_(*k*) for each *k* ∈ *K*). This motivates a straightforward model of how animals estimate value in partially observable environ-ments [6, 7], which we will refer to as the “Belief model”: First compute beliefs (Eqn. (6)), and then compute a linear transformation of those beliefs to a value estimate (Eqn. (7)), with weights obtained from TD learning.

### Learning state representations using recurrent neural networks

The Belief model assumes animals use a particular feature representation (i.e., beliefs) for estimating value. Here we ask what representations might emerge from solving the task of estimating value alone. It is useful to note that the Belief model can be written as follows:

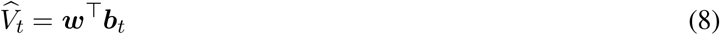

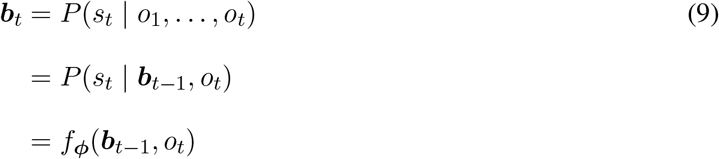

where *f* is a function parameterized by a specific choice of (fixed) parameters *ϕ*, and only ***w*** is learned. This latter equation has the same form as a generic recurrent neural network (RNN). This suggests a model could learn its own representation by treating *ϕ* as a learnable parameter. Taken together, this results in the following model for estimating value:

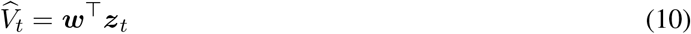

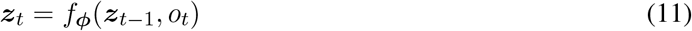

where ***z***_*t*_ ∈ ℝ^*H*^ is the activity of an RNN with *H* hidden units and parameters *ϕ*. Importantly, both *ϕ* and ***w*** can be learned simultaneously, using backpropagation through time to calculate Eqn. (4) with ***θ*** = [*ϕ* ***w***]. This allows the network to learn the representations, ***z***_*t*_, that are sufficient for estimating value. We will refer to this model as a “Value RNN” (see Materials and Methods).

Importantly, this RNN-based approach resolves all three challenges for learning a belief representation that we raised in the Introduction: 1) The model can learn from observations alone, as no information is provided about the statistics of the underlying environment; 2) the model’s size (parameterized by *H*, the number of hidden units) can be controlled separately from the number of states in the environment; and 3) the model’s only objective is to estimate value.

Though such a model has no explicit objective of learning beliefs, the network may discover a belief representation implicitly. We next asked what signatures, if any, would indicate the existence of a belief representation. In the sections that follow we develop an analytical approach for determining whether the Value RNN’s learned representations resemble beliefs.

### RNNs learn belief-like representations

As a working example, we will consider the probabilistic associative learning paradigm where dopamine RPEs were shown to be consistent with a belief representation [7, 13]. This has the added benefit of ensuring that the RNN-based approach described above can recapitulate these previous results.

This paradigm consisted of two tasks, which we will refer to as Task 1 and Task 2 (Fig 1). In both tasks, mice were trained to associate an odor cue with probabilistic delivery of a liquid reward 1.2-2.8s later. The tasks were each composed of two states: an intertrial interval (ITI), during which animals waited for an odor; and an interstimulus interval (ISI), during which animals waited for a reward. In Task 1, every trial contained both an odor and a reward. As a result, the animal’s observations could fully disambiguate the underlying state: An odor signaled a transition to the ISI state, while a reward signaled a transition to the ITI state. In Task 2, by contrast, reward on a given trial was omitted with 10% probability. This meant the underlying states were now only partially observable; for example, in Task 2 an odor signaled a transition to the ISI state with 90% probability.

**Fig 1.**
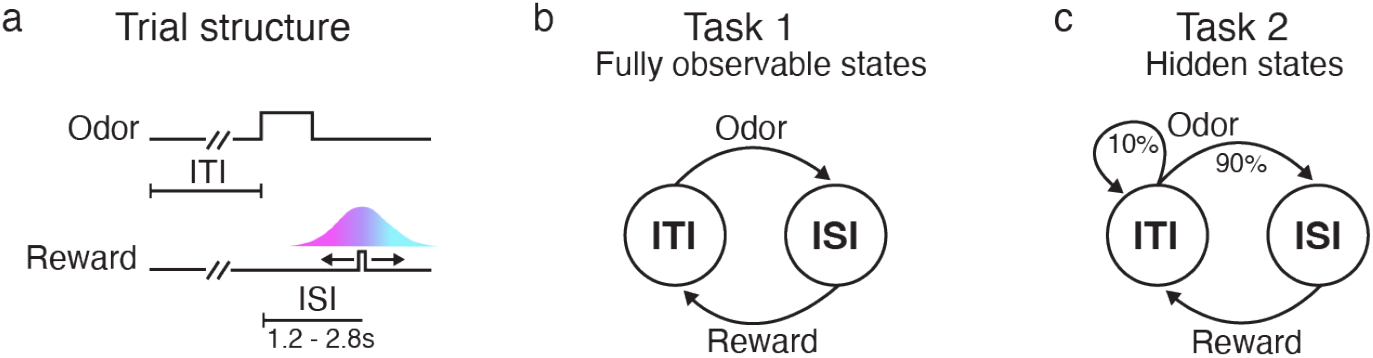
Associative learning tasks with probabilistic rewards and hidden states. **a**. Trial structure in Starkweather et al. [7, 13]. Each trial consisted of a variable delay (the intertrial interval, or ITI), followed by an odor, a second delay (the interstimulus interval, or ISI), and a potential subsequent reward. Reward times were sampled from a discretized Gaussian ranging from 1.2-2.8s (see Materials and Methods). **b-c**.. In both versions of the task, there were two underlying states: the ITI and the ISI. In Task 1, every trial was rewarded. As a result, an odor always indicated a transition from the ITI to the ISI, while a reward always indicated a transition from the ISI to the ITI. In Task 2, rewards were omitted on 10% of trials; as a result, an odor did not reveal whether or not the state transitioned to the ISI.

To formalize these tasks, we largely followed previous work [7, 13]. Each task was modeled as a discrete-time Markov process with states *s*_*t*_ ∈ {1, …, *K*}, where each *t* denotes a 200ms time bin (Fig 2A). These *K* “micro” states can be partitioned into those belonging to one of two “macro” states (corresponding to the ITI and the ISI; see Materials and Methods). At each point in time, the agent’s observation is one of *o*_*t*_ ∈ {*odor, reward, null*} (Fig 2B). For each task, we trained the Belief model, and a Value RNN (using a gated-recurrent unit cell [24], or GRU, with *H* = 50 hidden units), on a series of observations from that task to estimate value using TD learning (see Materials and Methods). Before training, the Value RNN’s representation consisted of transient responses to each observation (Fig S1). After training, we evaluated each model on a sequence of new trials from the same task (Fig 2C).

**Fig 2.**
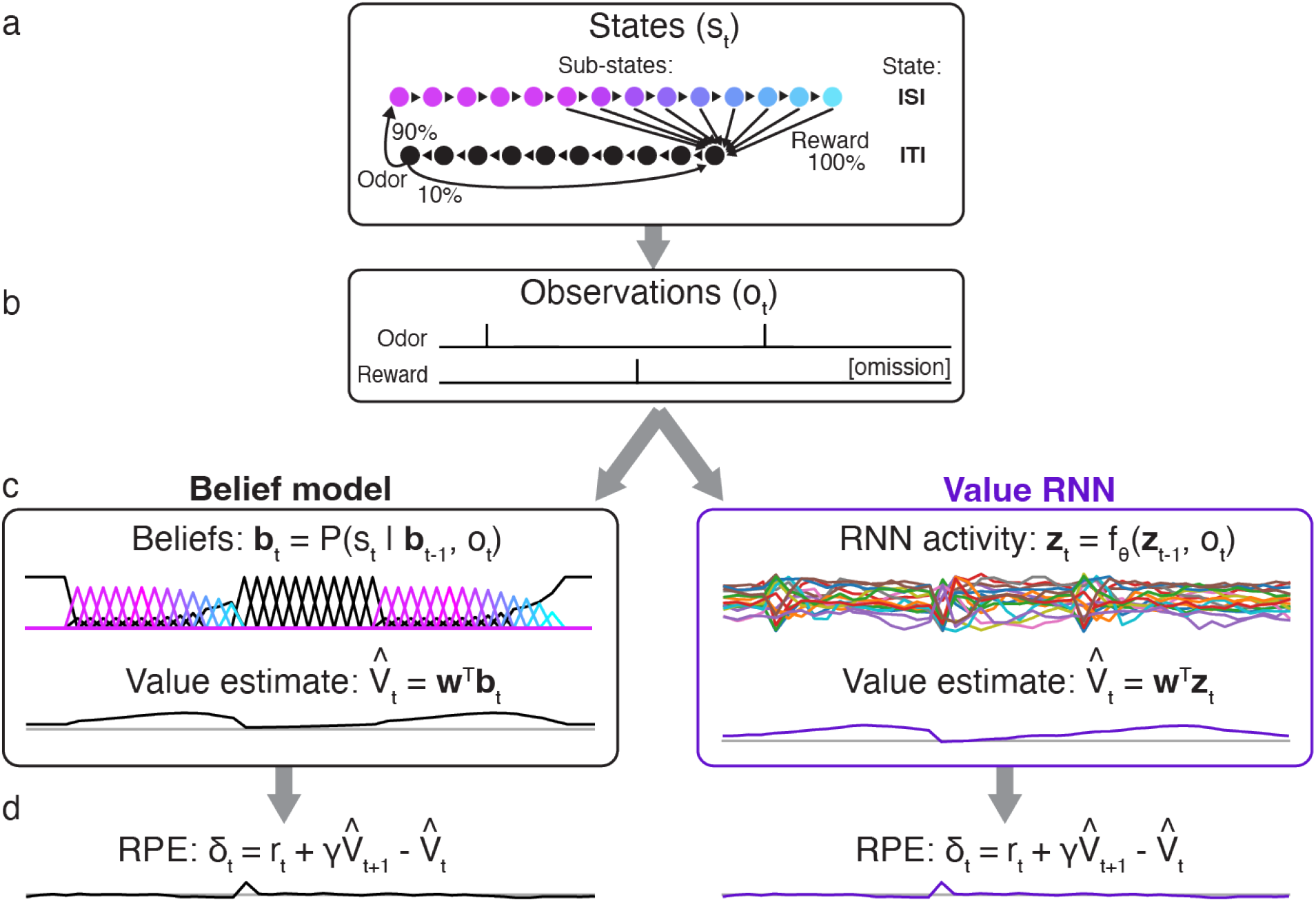
Observations, model representations, value estimates, and reward prediction errors (RPEs) during Task 2. **a**. State transitions and observation probabilities in Task 2. Each macro-state (ISI or ITI) is composed of micro-states denoting elapsed time; this allows for probabilistic reward times and minimum dwell times in the ISI and ITI, respectively. **b**. Observations emitted by Task 2 during two example trials. Note that omission trials are indicated only implicitly as the absence of a reward observation. **c**. Example representations (***b***_*t*_, ***z***_*t*_) and value estimates 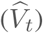 of two models (Belief model, left; Value RNN, right) for estimating value in partially observable environments, after training. **d**. After training, both models exhibit similar RPEs.

To confirm that this approach could recapitulate previous results, we measured the RPEs generated by each trained model (Fig 2D). Previous work showed that dopamine RPEs depended on the reward time differently in the two tasks, with RPEs increasing as a function of reward time in Task 1, but decreasing as a function of reward time in Task 2 [7] (Fig 3A). As in previous work, we found that this pattern was also exhibited by the Belief model (Fig 3B). We found that the RPEs of the Value RNN exhibited the same pattern (Fig 3C). In particular, the Value RNN’s RPEs became nearly identical to those of the Belief model after training (Fig 3D). We emphasize that the Value RNN was not trained to match the value estimate from the Belief model; rather, it was trained via TD learning using only observations. This result shows that, through training on observations alone, Value RNNs discovered a representation that was sufficient for both learning the value function as well as explaining empirical dopamine activity.

**Fig 3.**
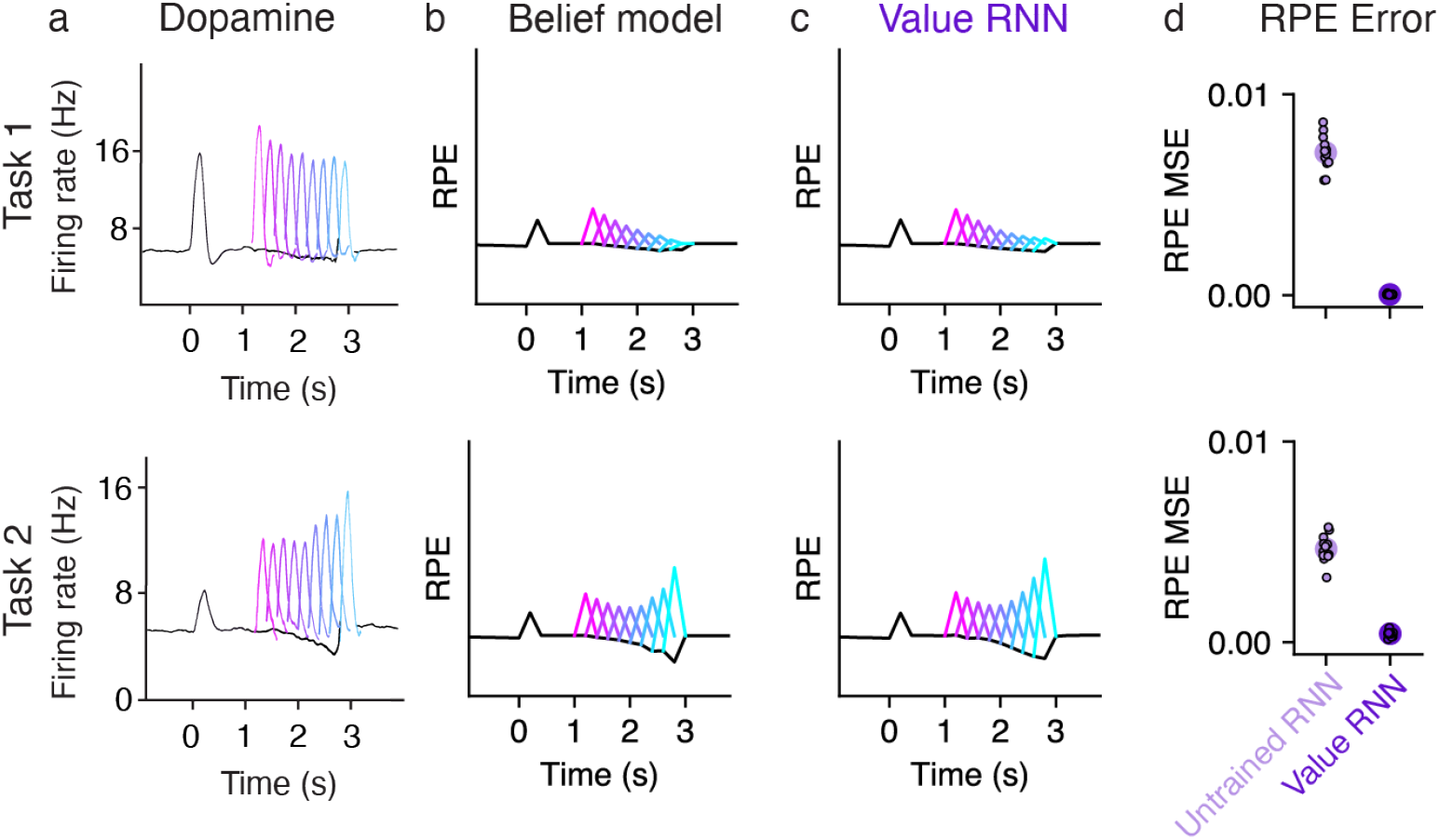
RPEs of the Value RNN resemble both mouse dopamine activity and the Belief model. **a**. Average phasic dopamine activity in the ventral tegmental area (VTA) recorded from mice trained in each task separately. Black traces indicate trial-averaged RPEs relative to an odor observated at time 0, prior to reward; colored traces indicate the RPEs following each of nine possible reward times. RPEs exhibit opposite dependence on reward time across tasks. Reproduced from Starkweather et al. [7]. **b-c**. Average RPEs of the Belief model and an example Value RNN, respectively. Same conventions as panel **a. d**. Mean squared error (MSE) between the RPEs of the Value RNN and Belief model, before and after training each Value RNN. Small dots depict the MSE of each of *N* = 12 Task 1 RNNs and *N* = 12 Task 2 RNNs, and circles depict the median across RNNs.

We next asked whether the Value RNN learned these tasks by forming representations that resembled beliefs. We considered three approaches to answering this question. First, we asked whether beliefs could be linearly decoded from the RNN’s activity. Next, because beliefs are the optimal estimate of the true state in the task, we asked whether RNN activity could similarly be used to decode the true state. Finally, we took a dynamical systems perspective, and asked whether the RNN and beliefs had similar dynamical structure.

### RNN activity readout was correlated with beliefs

We first asked whether the Value RNN’s activity was correlated with beliefs. Because the belief and RNN representations did not necessarily have the same dimensionality, we performed a multivariate linear regression to find the linear transformation of the RNN’s activity that came closest to matching the beliefs (see Materials and Methods). In other words, we found the linear transformation, 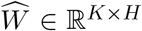, that could map the RNN’s activity, ***z***_*t*_ ∈ ℝ^*H*^, to best match the belief vector, ***b***_*t*_ ∈ ℝ^*K*^ :

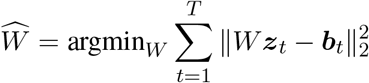

To quantify performance, we used held-out sessions to measure the total variance of the beliefs that were explained by the linear readout of RNN activity (*R*^2^; see Materials and Methods). We found that the Value RNN’s activity explained most of the variance of beliefs (Fig 4B; Task 1 *R*^2^: 0.69 ± 0.01, mean ± SE, *N* = 12; Task 2 *R*^2^: 0.76 ± 0.02, mean ± SE, *N* = 12), substantially above the performance when using the RNN activity before training (Task 1 *R*^2^: 0.47 ± 0.00, mean ± SE, *N* = 12; Task 2 *R*^2^: 0.50 ± 0.00, mean ± SE, *N* = 12). This is not a trivial result of the network’s training objective, as the Value RNN’s target (i.e., value) is only a 1-dimensional signal, whereas beliefs are a *K*-dimensional signal (here, *K* = 25). Nevertheless, we found that training the Value RNN to estimate value resulted in its representation becoming more belief-like, in the sense of encoding more information about beliefs.

**Fig 4.**
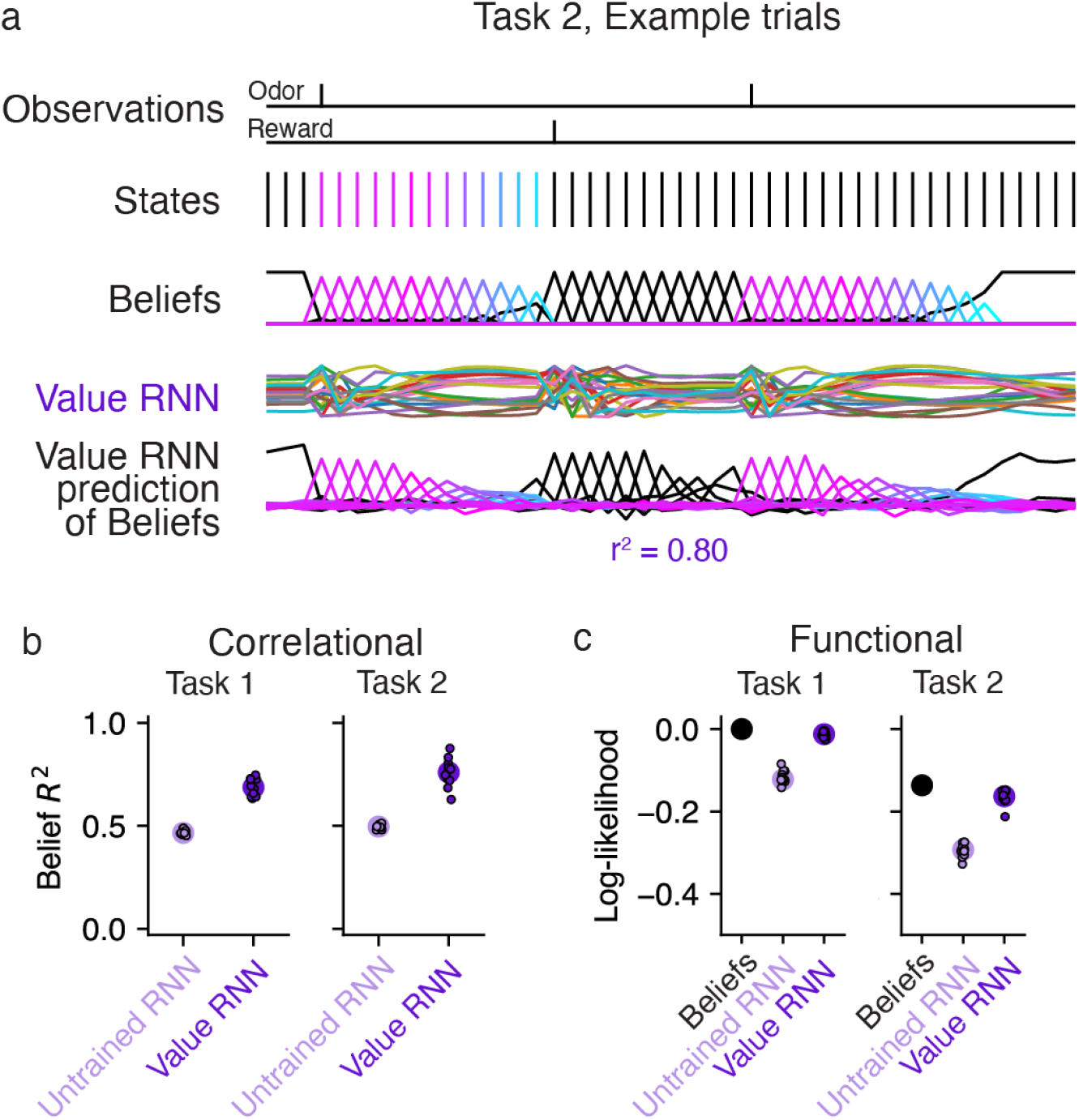
Value RNN activity readout was correlated with beliefs and could be used to decode hidden states. **a**. Example states, beliefs, and Value RNN activity from the same Task 2 trials shown in Fig 2. Note that the states following the second odor observation remain in the ITI (black) because the second trial is an omission trial. Bottom traces depict the linear transformation of the RNN activity that comes closest to matching the beliefs. Total variance explained (*R*^2^) is calculated on held-out trials. **b**. Total variance of beliefs explained (*R*^2^), on held-out trials, using different trained and untrained Value RNNs, in both tasks. Same conventions as Fig 3D). **c**. In purple, the cross-validated log-likelihood of linear decoders trained to estimate true states using RNN activity. Same conventions as Fig 3D). Black circle indicates the log-likelihood when using the beliefs as the decoded state estimate (i.e., no decoder is “trained”).

### RNN activity could be used to decode hidden states

The previous analysis assessed how much information about beliefs was encoded by RNN activity. Given that beliefs are distributions over hidden states, we asked whether the ground truth state could be decoded from the RNN’s activity. To do this, we performed a multinomial logistic regression to find a linear transformation of the RNN’s activity that maximized the log-likelihood of the true states (see Materials and Methods). We quantified performance on held-out sessions by evaluating the log-likelihood of the decoded estimates. Because the beliefs capture the posterior distribution of the state given the observations under the true generative model, the log-likelihood of the beliefs serves as a ceiling on performance. We found that the log-likelihoods of the decoders trained on the RNN’s activity approached those of the beliefs, and easily outperformed the decoders that used the activity of untrained RNNs (Fig 4C). Thus, training the RNN to estimate value resulted in a representation that could be used to more accurately decode the true state.

### RNN activity exhibited belief-like dynamics

One potential shortcoming of the above analyses is that we have not yet accounted for the dynamical nature of the belief representation: Belief updating can be thought of as a dynamical system describing how the posterior probability of each state evolves as a function of the observations. We therefore took a dynamical systems perspective [25, 26, 27] and asked whether the dynamics of RNN activity resembled the dynamics of the beliefs in each task.

We first asked whether beliefs and RNNs had similar fixed point structure, a standard approach to characterizing the computations performed by dynamical systems [25, 26, 27]. Here, a “fixed point” is a belief state that remains unchanged in the absence of any new observations. Specifically, we considered the fixed points of beliefs in the absence of observations (Fig 5A). In both tasks, the duration of the ITI is sampled from a geometric distribution, which has a constant hazard function. Thus, if the agent believes it is in the ITI (i.e., waiting for an odor), it should maintain this belief for as long as it receives no new observations (∅). Thus, the ITI belief state is a fixed point of the belief updates in both tasks (Fig 5A). Now consider when the agent is in the ISI (e.g., following an odor observation). In Task 2 (Fig 5A, bottom panel), the agent should maintain a nonzero belief in the ISI only for as long as there are possible reward times remaining—i.e., the first 2.8s, or 14 time steps—but after that point it should return to the ITI state. Thus, the ITI state is the only fixed point of the Task 2 beliefs. In Task 1, by contrast, there are no omission trials, and so the beliefs are simply undefined when there are no observations for more than 14 time steps. Nevertheless, for the purposes of characterizing the fixed points of beliefs, we can ask what an agent with Task 1 beliefs *could* do when faced with an omission trial. In this sense, an agent could maintain a belief in the ISI for any number of time steps *X >* 14, resulting in two fixed points when *X* → ∞, and one fixed point otherwise (Fig 5A, top panel). Thus, Task 1 beliefs can decay to the ITI fixed point at *any* point after 14 time steps, and may potentially have two fixed points.

**Fig 5.**
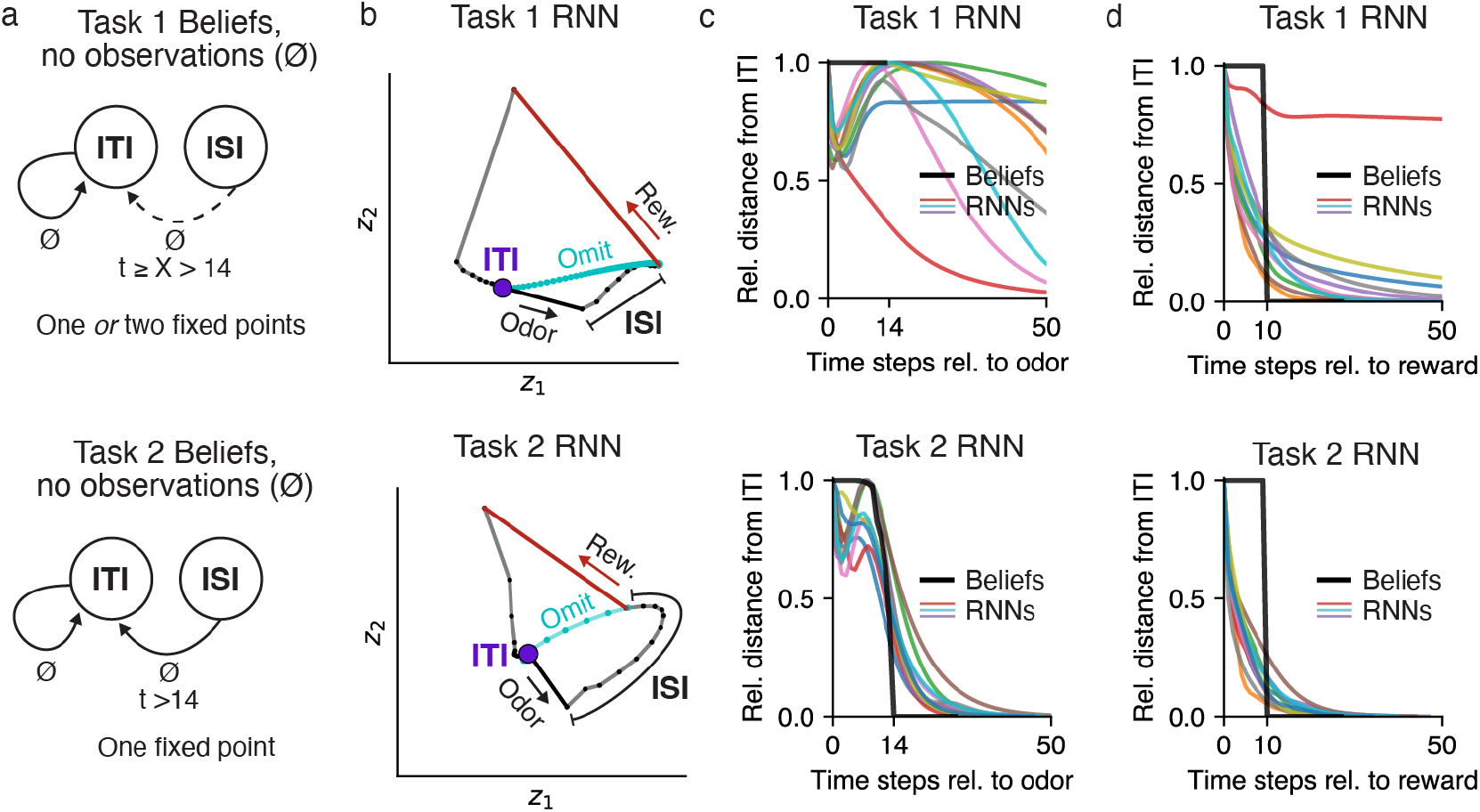
Value RNN dynamics resembled belief dynamics in each task. **a**. Dynamics of beliefs in Task 1 (top) and Task 2 (bottom). Black arrows indicate transitions between states in the absence of observations (∅) as a function of elapsed time, *t*, following an odor observation. ‘X’ indicates an unconstrained duration, and a dashed arrow indicates a transition that happens only when ‘X’ is finite. **b**. RNN activity at each time step (small black dots with connected lines) during an example trial in a 2D subspace identified using PCA. Putative ITI fixed point indicated as purple circle. Vectors indicate the response to odor (black) and reward (red). Activity during an omission trial is shown in cyan, though note that omission trials were present in training data only for Task 2. **c-d**. Average normalized distance of each model’s activity from its fixed point (identified numerically) following an odor (panel **c**) or reward (panel **d**) observation, over time. To allow comparing distances across models, each model’s distances were normalized by the maximum distance following each observation.

We asked whether the Value RNNs in each task exhibited similar dynamics. To build intuition, for each task we visualized the activity of a Value RNN on example trials (Fig 5B). To visualize the activity of the RNN’s 50 units, we used principal components analysis (PCA) to project the RNN’s activity into the top two dimensions that captured the most variance of the activity across trials; these two dimensions explained 83% and 79% of the total variance in the Task 1 and Task 2 Value RNN, respectively. We observed that each RNN’s activity was quite stable during the ITI (purple circle), suggestive of a fixed point. This activity was then perturbed by an odor observation (black vector), and continued to move through state space during the ISI. On rewarded trials, in response to a reward (red vector), RNN activity gradually returned to the same ITI location it started from (purple circle). We noted that the activity of both RNNs would also have converged to its original ITI location had the reward been omitted (cyan traces). Interestingly, this was true even for the Task 1 RNN, which did not experience omission trials during training. These visualizations suggested that these two example Value RNNs had a single fixed point (corresponding to the ITI), which we then confirmed numerically (see Materials and Methods). We next used the same numerical approach to identify the fixed points across all trained Value RNNs, and found similar results. In fact, only two Value RNNs had more than one fixed point; these were both Task 1 RNNs, which had a fixed point for both the ITI and the ISI. Overall, these analyses indicated that Value RNNs had a fixed point structure consistent with those of beliefs.

Despite the fact that most Value RNNs had a single fixed point regardless of which task they were trained on, we noted that the temporal dynamics of RNN activity differed across the two tasks following odor observations. For example, in the Task 2 RNN, following an odor observation, the activity moved gradually closer to the ITI state throughout the ISI (Fig 5B, bottom subpanel). These dynamics allowed the Task 2 RNN’s activity to return to the ITI state at the appropriate time on trials without reward (cyan trace). By contrast, in the Task 1 RNN, which did not experience trials without rewards during training, activity took much longer to return to the ITI. To quantify these differences, we initialized each RNN to its fixed point, provided an odor observation, and then measured the RNN’s activity over time in the absence of any subsequent reward. We then measured the distance of each RNN’s activity over time from its ITI fixed point (Fig 5C, colored traces).

We repeated this same analysis for beliefs (Fig 5C, black trace), allowing us to characterize the two models’ responses to an odor as a function of the distance of their representations from their ITI fixed point. We reasoned that, if the RNNs learned belief-like dynamics, the activity of Task 2 RNNs should return to the ITI as soon as possible after time step 14 (i.e., the largest reward time), which we found was largely the case (Fig 5C). By contrast, in Task 1, beliefs are undefined past time step 14 (because there are no omission trials), and so the activity of RNNs after this point was not constrained by the task. To quantify these differences across tasks, we calculated the time step at which each RNN’s activity returned within some threshold distance of its ITI fixed point. We refer to this quantity—which essentially measures ‘X’ in the top panel of Fig 5A—as the network’s *odor memory*. In fact, we found that *all* Task 1 RNNs had longer odor memories (310 ± 150, mean ± SE, *N* = 12) than Task 2 RNNs (49 ± 2, mean ± SE, *N* = 12). Overall, these features were consistent with the beliefs in the two tasks following an odor observation: beliefs in Task 2, but not Task 1, must quickly return to the ITI after the maximum possible reward time. We performed a similar analysis for reward observations, in which case the network activity in both tasks was expected to return to the ITI fixed point shortly after the minimum ITI duration (at time step 10). Here, we found that the activity of both Task 1 and Task 2 RNNs quickly returned to the ITI fixed point after this point (Fig 5D), again consistent with the beliefs in these tasks.

### RNNs with larger capacity had more belief-like representations

Thus far, we have considered the representations of Value RNNs with the same number of hidden units (*H* = 50). To understand whether the network’s size influences the types of representations learned, we next trained Value RNNs with a variable number of hidden units. We found that Value RNNs with as few as 2 hidden units could learn the value function, as evidenced by their RPEs matching those of the Belief model (Fig 6A). In other words, despite there being 25 discrete states in our implementation of this task (and, as such, beliefs were 25-dimensional), an RNN with a 2-dimensional representation was sufficient for performing the task. On the other hand, RNNs with fewer units had representations that were notably less belief-like, in terms of how well they linearly encoded beliefs (Fig 6B) and allowed for decoding the true state (Fig 6C). Thus, Value RNNs with 2 or more hidden units could all estimate value equally well, but only those with a sufficient number of hidden units featured representations that resembled beliefs.

**Fig 6.**
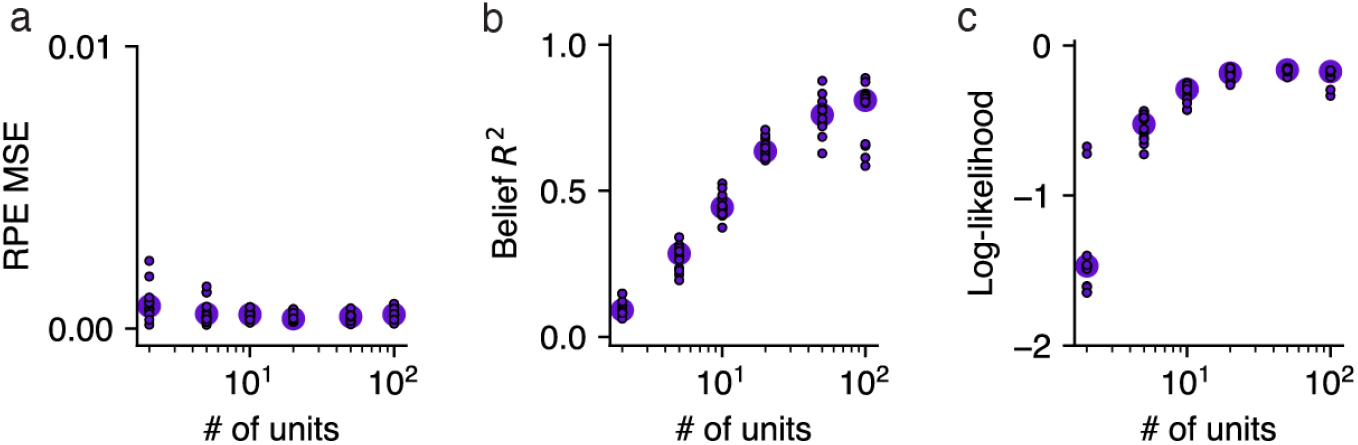
Value RNNs with larger capacity had more belief-like representations. **a**. Error between the Value RNN’s RPEs and those of the Belief model (“RPE MSE”; see Fig 3D) during Task 2, as a function of the number of units in the Value RNN. Each circle indicates the median across the *N* = 12 Value RNNs with the same number of units. **b**. Total variance explained (*R*^2^) of beliefs (see Fig 4B). Same conventions as panel **a. c**. Log-likelihood of the state decoder using Value RNN activity to estimate the true state (see Fig 4C). Dashed line indicates maximum possible log-likelihood (i.e., from Belief model). Same conventions as panel **a**.

## Generalization to other tasks

We showed that RNNs could be trained to estimate value in the tasks from Starkweather et al. [7], and that the representations of these RNNs became more belief-like as a result of training. We next assessed whether the same basic insights generalized to a different task, that of Babayan et al. [10]. In this task, similar to Task 1 of Starkweather et al. [7], each trial consisted of an odor followed by a deterministic reward. Unlike in the Starkweather task, in this task the reward quantity on each trial was varied in blocks. In Block 1, each trial consisted of a small (1 *μ*L) reward, while in Block 2 each trial consisted of a large (10 *μ*L) reward. As a result, we formalize the states in this task using pairs of ITI and ISI states, one for each block (Fig 7A). Importantly, the block identity was hidden to the animal, and was resampled uniformly every five trials. This meant that animals had to use the reward observations to infer which block they were currently in.

**Fig 7.**
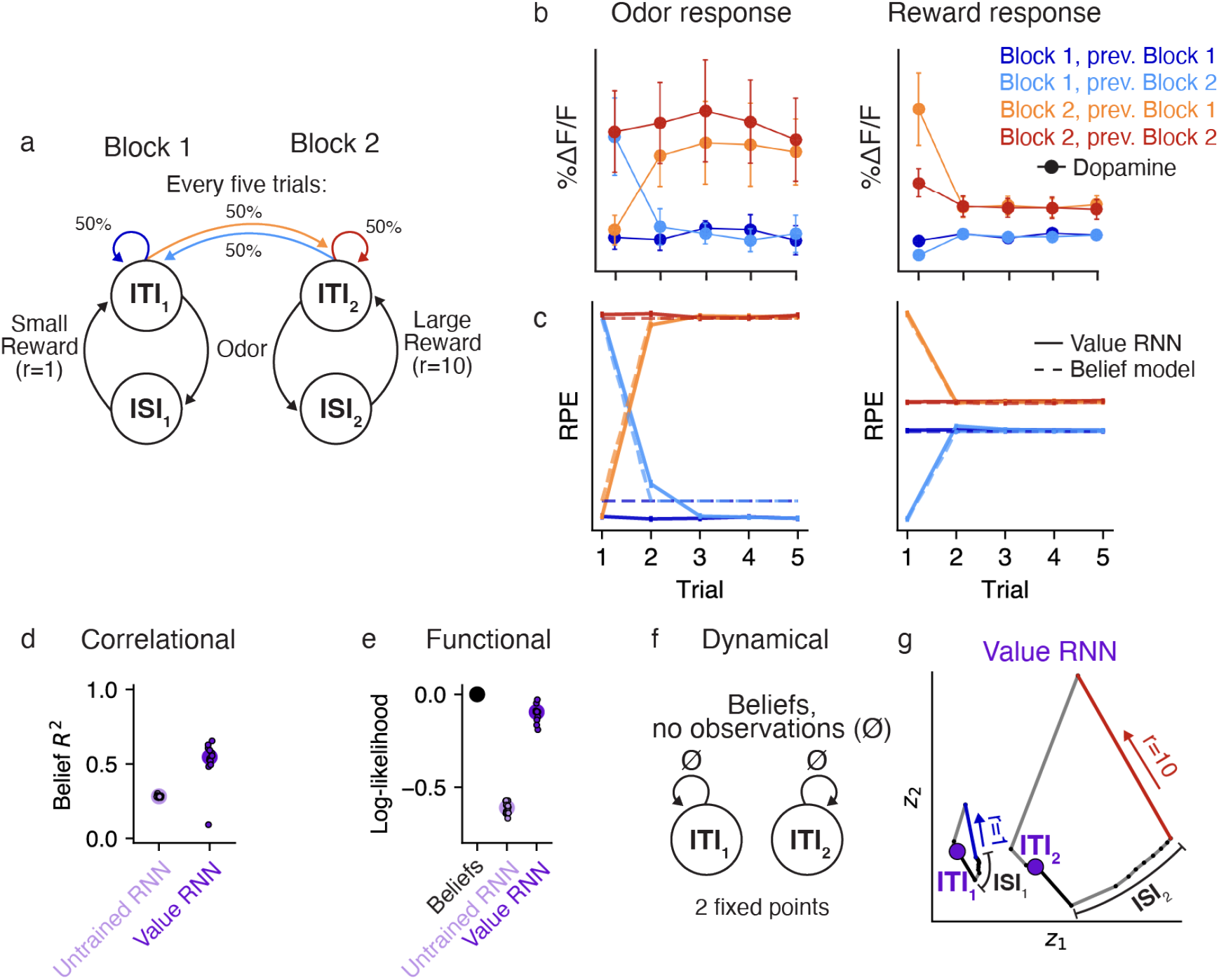
Value RNNs trained on Babayan et al. [10] reproduce Belief RPEs and learn belief-like representations. **a**. Task environment of Babayan et al. [10]. Each trial consists of an odor and a subsequent reward. The reward amount depends on the block identity, which is resampled uniformly every five trials (green). **b**. Average phasic dopamine activity in the VTA of mice trained on the task at the time of odor (left) and reward (right) delivery. Activity is shown separately as a function of the trial index within the block (x-axis) and the current/previous block identity (colors). Reproduced from Babayan et al. [10]. **c**. Average RPEs of the Belief model (dashed lines) and an example Value RNN (solid lines). Same conventions as panel **b. d**. Total variance of beliefs explained (*R*^2^) using a linear transformation of model activity. Same conventions as Fig 4B. **e**. Crossvalidated log-likelihood of linear decoders trained to estimate true states using RNN activity. Same conventions as Fig 4C. **f**. Dynamics of beliefs in the absence of observations. Same conventions as Fig 5A. **g**. Trajectories of an example Value RNN’s activity, in the 2D subspace identified using PCA, during an example trial from Block 1 (left) and Block 2 (right). These two dimensions explained 69% of the total variance in the Value RNN’s activity across trials. Putative ITI states indicated as purple circles. Same conventions as Fig 5B.

Previous work showed that the dopamine activity of animals trained on this task depended on the number of trials in the current block (Fig 7C), similar to what you would expect if animals used a belief representation (Fig 7D, dashed lines) [10]. To see if the Value RNN could reproduce these results, we trained *N* = 12 Value RNNs on this task. We found that Value RNNs exhibited nearly identical RPEs as the Belief model (Fig 7D). This was even true on probe sessions that included blocks with intermediate reward sizes, a setting in which both dopamine activity and belief RPEs exhibited a characteristic nonlinear relationship with reward size (Fig S2). These results indicate that the Value RNNs found a representation sufficient for estimating value despite the hidden states.

We next asked whether, as in the Starkweather task, the Value RNN’s representations resembled beliefs. To do this, we repeated the analyses in Fig 4. We found that the Value RNN’s activity could be linearly transformed to match the beliefs (Fig 7D), and that its activity could also be used to decode the hidden states in the task (Fig 7E), as compared to RNNs not trained on the task. We next took a dynamical systems approach, characterizing the fixed points of beliefs in this task. Similar to Task 1 of Starkweather et al. [7] (Fig 5A), beliefs in the present task should have a fixed point at the ITIs for Block 1 and Block 2 (Fig 7F). To assess whether this was the case in the Value RNN, we visualized an example RNN’s activity on the last few trials of each block, when the network should be confident as to the current block’s identity (given the reward observations on previous trials). During these trials, we observed two non-overlapping trajectories of activity for each block (Fig 7G). Following a reward observation, the RNN’s activity converged to a distinct location in state space corresponding to that block’s ITI. This suggested the RNN had two fixed points, as in the belief representation. In reality, these were not both truly fixed points, as the RNN’s activity did eventually return to a single fixed point given enough time without an observation (Fig S3A). However, the RNN’s two putative ITI states remained distinct across the range of ITI durations present in the training data (Fig S3B), allowing the network to keep these trajectories (and thus the states corresponding to each block) separate. These analyses suggest that Value RNNs trained on this task also exhibited belief-like representations.

### Untrained RNNs could also be used to estimate value and encode beliefs

In the sections above we analyzed Value RNNs whose representations had been trained through TD learning. Here we take an alternative approach, inspired by reservoir computing, and consider the representations of untrained RNNs. In reservoir computing, a static dynamical system, or “reservoir,” is combined with a learned linear readout. Given an appropriately initialized reservoir (e.g., an RNN), this approach can be used to approximate any nonlinear function [28]. Inspired by this approach, we explored whether we could choose a random initialization of our RNNs such that it was only necessary to learn a linear weighting of the RNN’s representation to form its value estimate (i.e., *ϕ* in Eqn. (11) was fixed throughout training). Because this model resembles an echo state network (“ESN”; a reservoir computer whose reservoir is an RNN [29]), we will refer to this model as a Value ESN.

We initialized each RNN by sampling the matrix of recurrent weights as a random orthogonal matrix scaled by a single gain parameter [30], an approach commonly used to initialize RNNs (see Materials and Methods). The gain effectively modulated the duration of the network’s transient response to inputs (Fig 8A-B), such that larger gains led to larger odor memories (Fig 8C). In agreement with previous work, we found that when the gain was above a critical value (“the edge of chaos” [30]), the network’s activity never decayed back to baseline (e.g., gain *>* 2 in Fig 8C).

**Fig 8.**
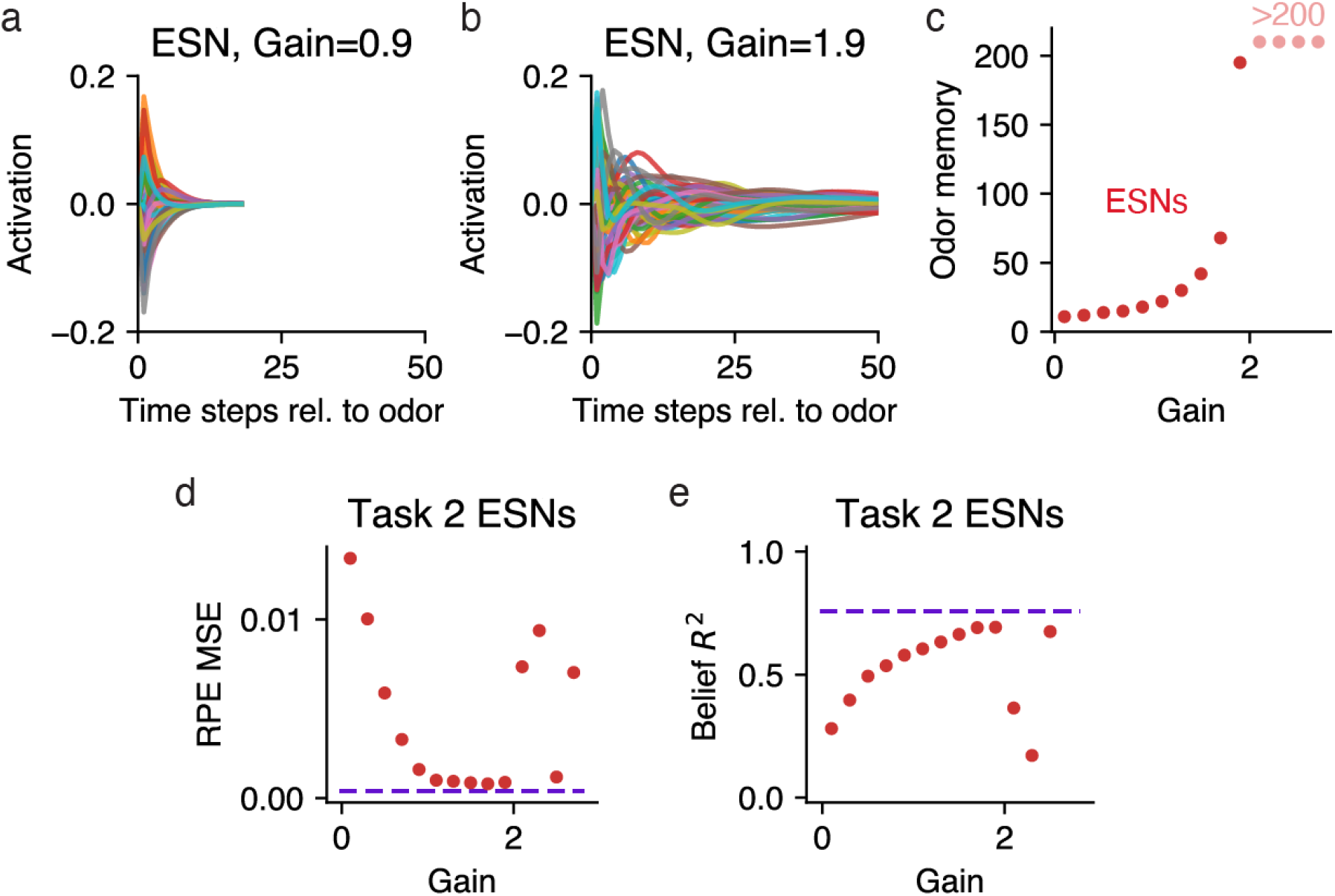
Untrained RNNs can also be used to estimate value. **a**. Time-varying activations of 20 example units in response to an odor input, in an untrained RNN (“ESN”) with 50 units, initialized with a gain of 0.1. **b**. Same as panel **a**, but for an initialization gain of 1.9, and a wider range shown on the x-axis. **c**. Number of time steps it took each ESN’s activity to return to its fixed point following an odor observation (“odor memory”; see Materials and Methods), as a function of ESN initialization gain. Points labeled “*>*200” indicate those that did not fit within the plot. **d**. RPE MSE (see Fig 3D) as a function of ESN initialization gain, after training each ESN’s value weights to estimate value during Task 2. Dashed line indicates average levels of a Task 2 Value RNN with the same number of units. **e**. Belief *R*^2^ (see Fig 4B) as a function of ESN initialization gain. Same conventions as panel **d**.

To see if Value ESNs could be trained to estimate value, we trained Value ESNs on Starkweather Task 2, varying the gain for each network. During training, only the network’s value weights were modified to estimate the value function. We found that for a range of different gains, the resulting Value ESN could estimate value nearly as well as Value RNNs, and recapitulate the experimentally observed dopamine patterns (Fig 8D). Interestingly, the Value ESN’s representations could even be linearly transformed to match beliefs (Fig 8E). We emphasize that the Value ESN’s representation was determined solely by its initialization; given an appropriate initialization, the Value ESN’s representation could effectively act as a set of temporal basis functions, allowing the network to match any downstream target, including beliefs. However, in terms of dynamics, the Value ESN’s dynamics differed substantially from those of the beliefs: For the best performing Value ESN (with a gain of 1.9), following an odor observation, the RNN’s activity returned to its fixed point after around 200 time steps (Fig 8C), whereas Task 2 beliefs return to the ITI point in 15 time steps. These results show that allowing the RNN to modify its representation during training led to its representation becoming even more belief-like than expected from a more carefully initialized RNN.

## Discussion

We have shown that training RNNs to estimate value in partially observable environments yields representations that resemble beliefs. Specifically, we showed that, after training, the RNN activity i) could be linearly transformed to approximate beliefs, ii) could be used to decode the true state in the environment, and iii) had a dynamical structure consistent with beliefs.

From a theoretical perspective, using an RNN resolves the problem of how to compute belief states, by replacing the fine-tuned Bayesian machinery needed for beliefs with a more general learned function approximator (e.g., a recurrent neural network). Our results illustrate that, to perform tasks in partially observable environments, it is not necessary for an agent or animal to explicitly estimate states using a belief representation; rather, agents can learn a sufficient representation for solving the task from observations alone. This is promising from a normative perspective, as it shows how neural circuits might come to compute theoretical features such as beliefs without that objective needing to be explicitly learned. Moreover, there is a growing toolkit for reverse engineering RNN solutions [26, 27, 31], which can shed light on learned mechanisms of value computation.

One potential benefit of the Value RNN over the Belief model for estimating value is the ability to separate the capacity of the model (i.e., the number of hidden units in the RNN) from the size of the state space in the environment. As we showed in Fig 6, Value RNNs with fewer units discovered a representation that was more compressed than beliefs, but nevertheless sufficient for performing the task at hand. Value RNNs with more units had more belief-like representations. This suggests a potentially useful trade-off, in which agents could choose to allocate more capacity to a task in exchange for more belief-like representations. Such a trade-off may be a relevant feature for biological organisms, who must be able to perform tasks such as value estimation in complex environments where it may not always be feasible to learn the full belief representation.

From a methodological perspective, this work can serve as a blueprint for how to bridge analyses of neural computation across levels of abstraction. In future work, we hope to apply this framework to test neural models of how animals perform associative learning via reinforcement learning. For instance, previous work has suggested that prefrontal cortex may perform state estimation in tasks with hidden states [6, 12, 7, 13]. The same tools we apply here to artificial neural networks can also be applied to neural activity recorded from animals performing the same task. For instance, if cortex implements something like a Value RNN, cortical activity may show a longer transient response to odors during Task 1 than in Task 2 (Fig 5). On the other hand, if activity is more like a Value ESN, cortical representations should be largely the same in both tasks.

Traditional models of how animals perform trace conditioning tasks like the ones we consider here make a variety of implicit assumptions about how animals represent the passage of time [32, 33]. For example, the state space shown in Fig 2A, which forms the basis of the belief representation, conceptualizes the passage of time in the form of microstates. Many modeling efforts require even more assumptions to account for scalar variability in animals’ estimates of elapsed time, such as by incorporating a more complex set of temporal basis functions. In our model, by contrast, the Value RNN’s time-varying representation of inputs is learned through training. We observed that individual units in our RNNs were tuned to elapsed time relative to observations, and that the temporal precision of tuning decreased with elapsed time (Fig S1), both of which are standard assumptions of microstate representations [34]. Similarly tuned “time cells” have been observed in the striatum [35], hippocampus [36], and prefrontal cortex [37]. Our modeling suggests that at least some assumptions about microstate representations may be redundant in the sense that they may emerge naturally in recurrent networks that are trained to perform reinforcement learning. This viewpoint resonates with the idea (reviewed in [38]) that delay encoding can arise as an emergent property of neural network dynamics.

Previous work has shown that, in animals, prefrontal cortex activity is a necessary component of animals’ state representations [13]. This work found that inactivating prefrontal cortex in the Starkweather task led to animals’ RPEs in Task 2 resembling the RPEs of Task 1. This is what one would expect if prefrontal cortex was involved in estimating a belief in omission trials. In fact, both Tasks 1 and 2 include another form of uncertainty, which is the reward time on each trial. The fact that prefrontal inactivation did *not* interact with animals’ estimates of timing suggests that different neural circuits may form belief-like representations specific to particular types of state uncertainty (e.g., temporal uncertainty versus reward uncertainty). In the present work, our RNNs should be thought of as a generic computational model, and not a model of individual brain regions. These networks had a generic architecture and only a single layer; as a result, our model would be unable to distinguish between different sources of uncertainty. Nevertheless, it is an interesting question how architectural considerations, such as layer connectivity and the dominance of feedforward versus recurrent connectivity, might contribute to where in the brain different belief-like representations are formed.

Here we have shown that computing beliefs explicitly is not a necessary precursor for optimally performing a task in partially observable environments. Nor is it required to reproduce experimentally observed patterns of dopamine neuron activity. Nevertheless, beliefs are an efficient representation in that they are sufficient for solving *any* task in the same environment. Thus, beliefs may be a desirable representation for animals, who may need to achieve a range of different goals in the same environment (e.g., finding water when thirsty, but finding food when hungry) without having to learn a representation in each of these tasks separately. Future work should explore whether a dedicated belief mechanism is necessary in these multi-task settings, or if the RNN framework we present here can also yield representations that effectively generalize to new tasks in the same environment.

## Materials and Methods

### Task implementation

In each experiment, at each time step *t*, agents received two observations: an odor cue, *c*_*t*_ ∈ {0, 1}; and a reward, *r*_*t*_ ∈ {0, *r*}, where *r >* 0 depended on the task (see below). We will refer to the total observation vector as ***o***_*t*_ = [*c*_*t*_ *r*_*t*_]. We treated each time step as equal to 200ms.

Each trial began with an intertrial interval (ITI), *t*_*IT I*_ ∈ N, during which there were no observations (***o***_*t*_ = 0 for *t < t*_*IT I*_). The ITI (offset by a minimum delay of 10 time steps) was sampled as *t*_*IT I*_ − 10 ∼ Geom 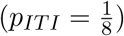, where Geom(*p*) is a geometric distribution with parameter *p*.

Following the ITI, a single odor cue was presented as *c*_*t*_ = 1 for *t* = *t*_*IT I*_. The cue was then followed by another interval with no observations, called the interstimulus interval (ISI), *t*_*ISI*_ ∈ 𝒩. A reward was then presented as *r*_*t*_ *>* 0 for *t* = *t*_*IT I*_ + *t*_*ISI*_, after which point the trial terminated. The details of the ISI and reward size depended on the specific task, as described below.

### Starkweather tasks

There were two versions of this task. In both Tasks 1 and 2, every non-zero reward size had *r*_*t*_ = 1. In Task 2, with probability *p*_*omission*_ = 0.1, the reward on a given trial was omitted, such that *r*_*t*_ = 0 for the duration of the trial. In both tasks, the ISIs on each trial were sampled from a discretized Gaussian with mean *μ* = 10, standard deviation *σ* = 2.5, and range 6 ≤ *t*_*ISI*_ ≤ 14, as in Starkweather et al. [7].

### Babayan task

In this task, non-zero reward sizes were determined in blocks of trials. In block 1, the non-zero reward size was *r*_*t*_ = 1, while in block 2 the non-zero reward size was *r*_*t*_ = 10. Each block consisted of 5 sequential trials. Block identities were sampled uniformly with equal probability. On all trials, the ISIs were uniformly sampled as *t*_*ISI*_ ∼ Unif({9, 10, 11}). For Fig S2, sessions also included blocks of intermediate rewards: *r*_*t*_ ∈ {1, 2, 4, 6, 8, 10}, where block identities were sampled in similar proportions to those used in Babayan et al. [10] (i.e., blocks with *r*_*t*_ = 1 or *r*_*t*_ = 10 comprised ∼90% of the total trials).

### Recurrent neural network implementation

We trained recurrent network models, in PyTorch, on multiple tasks to estimate value. Each *Value RNN* consisted of a GRU cell [24] with *H* ∈ ℕ units, followed by a linear readout of value. At each time step *t*, the RNN received observations, ***o***_*t*_ ∈ ℝ^2^, from a given experiment. The RNN’s representation can be written as ***z***_*t*_ = *f*_***ϕ***_(***o***_*t*_, ***z***_*t*−1_) given parameters *ϕ*. The RNN’s output was the value estimate 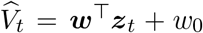, for ***w*** ∈ ℝ^*H*^ and *w*_0_ ∈ ℝ. The full parameter vector ***θ*** = [*ϕ* ***w*** *w*_0_] was learned using TD learning. This involved backpropagating the gradient of the squared error loss 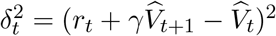 with respect to 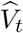 on episodes composed of 20 (Starkweather task) or 50 (Babayan task) concatenated trials. Unless otherwise noted we used *γ* = 0.93 as in Starkweather et al. [13].

For each task, and for each *H* ∈ {2, 5, 10, 20, 50, 100} units, we trained *N* = 12 networks. Prior to training each network, the weights and biases of the GRU (i.e., *ϕ*) were initialized using PyTorch’s default of 𝒰 (−*a, a*) where 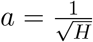. Training then proceeded for a maximum of 150 epochs on a session of 10,000 trials, with a batch size of 12 episodes. Training was stopped early if the loss increased for 4 consecutive epochs. Gradient updates used Adam with an initial learning rate of 0.003. No hyperparameter search was performed to fine-tune these choices. In the text, we refer to *Value RNNs* as the result of this training process, while *Untrained RNNs* are those that have been similarly initialized but not trained.

The *Value ESN* was similar to a Value RNN, except that it was initialized differently, and *ϕ* was frozen during training (i.e., the only learned parameters were ***w*** and *w*_0_). For initialization, we did the following (“Tensorflow-style” initialization). All of the GRU’s biases were initialized to zero. The GRU’s recurrent weights were sampled as a random orthogonal matrix using `torch.nn.init.orthogonal_` with a given gain [30]. The GRU’s input weights were sampled as 𝒰 (−*a, a*) where 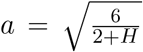, using `torch.nn.init.xavier_uniform_` [39].

### State and belief representations

Given a task with hidden states *s* ∈ {1, …, *K*}, the belief, ***b***_*t*_ ∈ [0, 1]^*K*^, is the posterior probability distribution over each possible state [17]. The tasks we analyze here are technically discrete-time semi-Markov processes, and so we follow previous work in formulating them equivalently as Markov processes with micro-states defined over each relevant discrete time step [6, 7]. In this setting, observations occur at the transition between states. As a result, the belief in state can be written as:

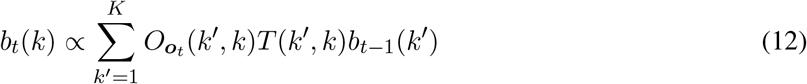

where *T* ∈ [0, 1]^*K*×*K*^ is the matrix of transition probabilities, and *O*_*o*_(*k*^′^, *k*) is the probability of observing *o* after making a transition from *k* to *k*^′^.

### Starkweather tasks

In both Tasks 1 and 2 there are three distinct observations, ***o***_*t*_ ∈ {[0 0], [1 0], [0 *x*]}, which we refer to as the null, odor, and reward observations, respectively. Let the possible reward times be *t*_*ISIs*_ = {6, …, 14}. The maximum reward time is max(*t*_*ISIs*_) = 14, and so we let states 1 − 14 be the ISI microstates. The ITI is a Geometric distribution plus a minimum ITI of *t*_*IT I*_ = 10, and so we let states 15 − 25 be the ITI microstates. There are *K* = 25 total states.

We first define the observation probabilities, *O*_*null*_, *O*_*odor*_, *O*_*reward*_ ∈ {0, 1}^*K*×*K*^, where *O*_*o*_(*k*^′^, *k*) indicates the probability of having transitioned from state *k* to state *k*^′^ upon observing *o* ∈ {*null, odor, reward*}. Each *O*_*o*_(*k*^′^, *k*) = 0 except at the following:

- *O*_*null*_(*t* + 1, *t*) = 1 for all *t* ≠max(*t*_*ISIs*_)
- *O*_*null*_(*K, K*) = 1
- *O*_*odor*_(*t, K*) = 1 for *t* = 1 (Task 1) or *t* ∈ {1, max(*t*_*ISIs*_) + 1} (Task 2)
- *O*_*reward*_(max(*t*_*ISIs*_) + 1, *t*) = 1 for *t* ∈ *t*_*ISIs*_

To define the transition probabilities, let *p*_*t*_ ∈ [0, 1] be the probability of receiving reward at time *t* ∈ *t*_*ISIs*_, *F*_*t*_ = Σ _*t′*≤*t*_ *pt′* the cumulative probability, and *h*_*t*_ = *p*_*t*_*/*(1 − *F*_*t*_) the hazard. Recall that 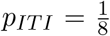. Then *T* (*k*^′^, *k*) = 0 except at the following:

- *T* (*t* + 1, *t*) = 1 for *t* ∉ max(*t*_*ISIs*_)
- *T* (*t* + 1, *t*) = 1 − *h*_*t*_ and *T* (max(*t*_*ISIs*_) + 1, *t*) = *h*_*t*_ for *t* ∈ *t*_*ISIs*_
- *T* (*K, K*) = 1 − *p*_*IT I*_
- *T* (max(*t*_*ISIs*_) + 1, *K*) = *p*_*IT I*_*p*_*omission*_
- *T* (1, *K*) = *p*_*ITI*_ (1 − *p*_*omission*_)

### Babayan task

The states in this task can be thought of as two copies of the beliefs in the Starkweather tasks, with one copy for each of the two blocks. (Note that *t*_*ISIs*_ = {9, 10, 11}, *K* = 22, and the hazard probabilities must be modified from the Starkweather task to account for the different reward timing distribution.) Each copy has 11 ISI microstates (because the maximum reward time is max(*t*_*ISIs*_) = 11) and 11 ITI microstates. Let 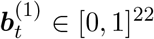 and 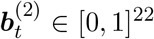 be the beliefs for the substates of block 1 and block 2, respectively. Then the full belief ***b***_*t*_ ∈ [0, 1]^44^ is as follows:

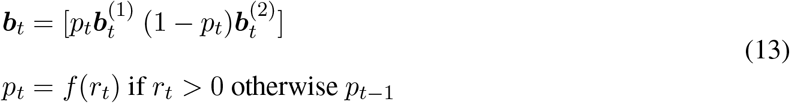

where *p*_*t*_ ∈ [0, 1] is the estimated probability of being in block 1, and *f* is our likelihood function mapping nonzero rewards, *r*_*t*_, to the estimated probability of being in block 1 vs. block 2. In other words, we modeled the belief in the block identity as being a function only of the most recently observed reward. We defined *f* following the original paper: Let *ϕ*(*r*_*t*_; *μ, σ*) be the pdf of a Normal distribution with mean *μ* and standard deviation *σ*_*r*_ *>* 0. Then 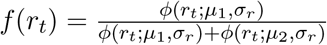, where *μ*_1_ = 1 and *μ*_2_ = 10 are the rewards amounts in block 1 and 2, respectively. Here we assumed *σ*_*r*_ was arbitrarily small, so we used *σ*_*r*_ = 0.001.

## Model analyses

We analyzed exemplars from each model class (Beliefs, Value RNNs, Untrained RNNs, Value ESNs) using two sessions of 1,000 concatenated trials each, using the same task parameters as those used when training the RNNs (see above). The first session was used for fitting any parameters relevant to the analysis (i.e., value weights, regression weights, decoding weights), while the second session was used for evaluation. Because the RNN’s responses were deterministic functions of their inputs, prior to analysis we added noise to all RNN representations to prevent overfitting during regression and decoding as follows. Let *σ*_*i*_ *>* 0 be the sample standard deviation of the activity of hidden unit *i* across trials. Then we added zero-mean Gaussian noise with a standard deviation of 0.01*σ*_*i*_ to this unit’s activity, so that each unit had the same SNR: 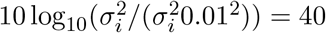dB.

### Value estimates

Each model’s value estimate was given by 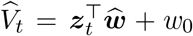, where 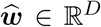 and *w*_0_ ∈ ℝare the value weights, and ***z***_*t*_ ∈ ℝ^*D*^ is the model’s representation at time *t*. (For the belief model, ***z***_*t*_ = ***b***_*t*_.)

We estimated the value weights, 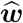, using Least-Squares TD (LSTD) [22]: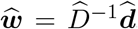, where 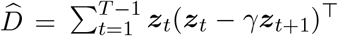 and 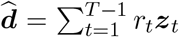. To ensure each model’s value function was estimated using the same procedure, we used LSTD even for the models including RNNs, even though the Value RNNs and Value ESNs learned their own value weights during training.

### Reward prediction errors

To assess how close each RNN’s RPEs came to the RPEs found using the belief model, we defined an RNN’s RPE error using the mean squared error: 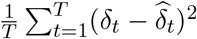, where 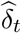 is the RNN’s RPE, and *δ*_*t*_ is the RPE from the belief model. Because each trial had at most one reward delivery, for simplicity we considered the RPEs only at the time of reward delivery on each trial (i.e., the *t* in the above equation refers to a trial and not a time step); this simplification did not affect our results.

### Belief *R*^2^

To assess how much variance of the beliefs could be explained by each model’s learned representation, we used multivariate linear regression:

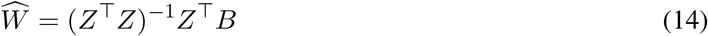

where *Z* ∈ ℝ^*T* ×(*H*+1)^ is the matrix whose *t*^*th*^ row is [***z***_*t*_ 1], *B* ∈ ℝ^*T* ×*K*^ is the matrix whose *t*^*th*^ row is ***b***_*t*_, and 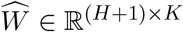.

To evaluate model fit, let Var 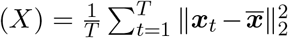, where ***x***_*t*_ is the *t*^*th*^ row of *X*, and 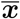 is the mean of the rows of *X*. Then we calculated the total variance explained:

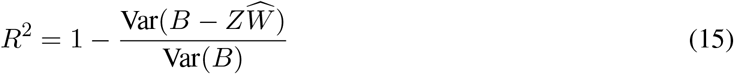

### State decoding

We asked where we could find a decoder that could infer the underlying state, *s*_*t*_ ∈ {1, …, *K*}, using an affine transformation of the RNN’s representation, ***z***_*t*_ ∈ ℝ^*H*^. To find such a decoder, we first standardized each model representation (considering each dimension of ***z***_*t*_ in isolation) to have zero mean and unit variance. We then performed a multinomial logistic regression using scikit-learn’s `linear model.LogisticRegression` with the parameters `multi class=“multinomial”`, `C=1`, and `max iter=1e4`.

After training, the decoder’s estimated state probabilities over *s*_*t*_ are:

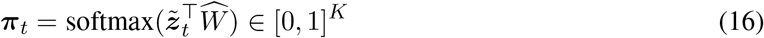

where 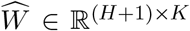 contains the decoder parameters; 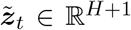 is the model representation at time *t* after standardization, plus an extra constant column of 1’s to fit the offset; and the softmax function normalizes the vector to be a valid probability over the *K* values of *s*_*t*_.

To evaluate the resulting decoder, we calculated the model’s log-likelihood (*ℓ*) on the evaluation session as follows:

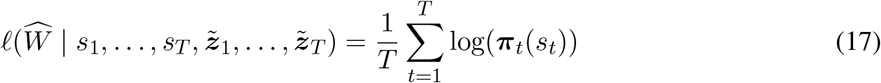

where ***π***_*t*_(*s*_*t*_) ∈ [0, 1] is the 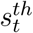 entry of the vector ***π***_*t*_. We calculated the log-likelihood for the belief model similarly, except instead of training a decoder we used ***π***_*t*_ = ***b***_*t*_. For the Babayan task (Fig 5E), we calculated the log-likelihood on all trials except the first trial in each block. This was necessary for the beliefs to act as an upper-bound on the log-likelihood, because we defined the beliefs in a way that did not assume knowledge of the number of trials in each block.

### Dynamics analysis

An RNN with parameters *ϕ* has a hidden state that evolves as ***z***_*t*_ = *f*_*ϕ*_(***o***_*t*_, ***z***_*t*−1_). Conditioned on a particular constant input, ***o***, an RNN is at a *fixed point* when 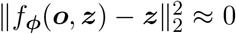. Numerically, we can simply look for ***z*** where 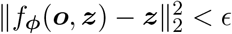. For our analyses below we chose *ϵ* = 1 × 10^−5^.

#### Identifying fixed points (Fig 5B, Fig 7G)

During training, RNNs received three distinct types of inputs, ***o***_*t*_ ∈ {[0 0], [1 0], [0 *r*]}, which we refer to as the null (∅), odor, and reward inputs, respectively. Under the beliefs of the Starkweather and Babayan tasks, the odor and reward inputs always result in a change in the beliefs. As a result, any fixed points of the beliefs must be conditional on the null input, ∅. We therefore sought to identify an RNN’s fixed points conditional on a null input. To do this, we took a numerical approach: We initialized the RNN to a random state, applied the null input until the RNN’s activity converged, and then repeated this process across different random states to get a candidate set of fixed points. More precisely, we considered *N* = 20 randomly selected values of ***z*** in the testing data following an odor or reward observation as a set of starting seeds. For each starting seed, ***z***_0_, we computed the RNN’s representation, ***z***_*t*_, over time, given no further odor or reward observations: ***z***_*t*_ = *f*_***ϕ***_(∅, ***z***_*t*−1_). We repeated this process until 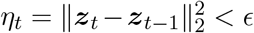. We then added ***z***_*t*_ to our list of candidate fixed points, ℱ. For each pair of candidate fixed points within a distance 1 × 10^−3^ of each other, we considered these to be the same fixed point.

#### Odor and reward memory duration (Fig 5C, Fig 5D)

For each Value RNN with a single fixed point, we measured the network’s odor (or reward) memory as follows. We initialized each RNN to its fixed point, ***z***_0_, and then provided an odor (or reward) observation at time *t* = 1. We then measured the RNN’s representation, ***z***_*t*_, over time, given no further odor or reward observations: ***z***_*t*_ = *f*_*ϕ*_(∅, ***z***_*t*−1_), for *t >* 1. For each *t*, we calculated the distance of the activity from its fixed point: 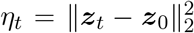 (Fig 5C). We repeated this until *η*_*t*_ converged to zero, defining the *odor memory* (or *reward memory*) as the *t* at which *η*_*t*_ *<* 1 × 10^−3^ (Fig 5D).

## Data Availability Statement

All data and code used for analysis and plotting is available on a GitHub repository at https://github.com/mobeets/value-rnn-beliefs/

## Author Contributions

**Conceptualization:** Jay A. Hennig, Sandra A. Romero Pinto, Takahiro Yamaguchi, Naoshige Uchida, Samuel J. Gershman

**Formal analysis:** Jay A. Hennig

**Investigation:** Jay A. Hennig

**Funding acquisition:** Samuel J. Gershman, Naoshige Uchida, Scott W. Linderman

**Methodology:** Jay A. Hennig, Samuel J. Gershman

**Supervision:** Scott W. Linderman, Naoshige Uchida, Samuel J. Gershman

**Visualization:** Jay A. Hennig

**Writing - original draft:** Jay A. Hennig

**Writing - review & editing:** Jay A. Hennig, Sandra A. Romero Pinto, Takahiro Yamaguchi, Scott W. Linderman, Naoshige Uchida, Samuel J. Gershman

## Acknowledgments

This work was funded by NIH U19 NS113201-01 and Air Force Office of Scientific Research grant FA9550-20-1-0413.

## Supplemental Figures

**Fig S1.**
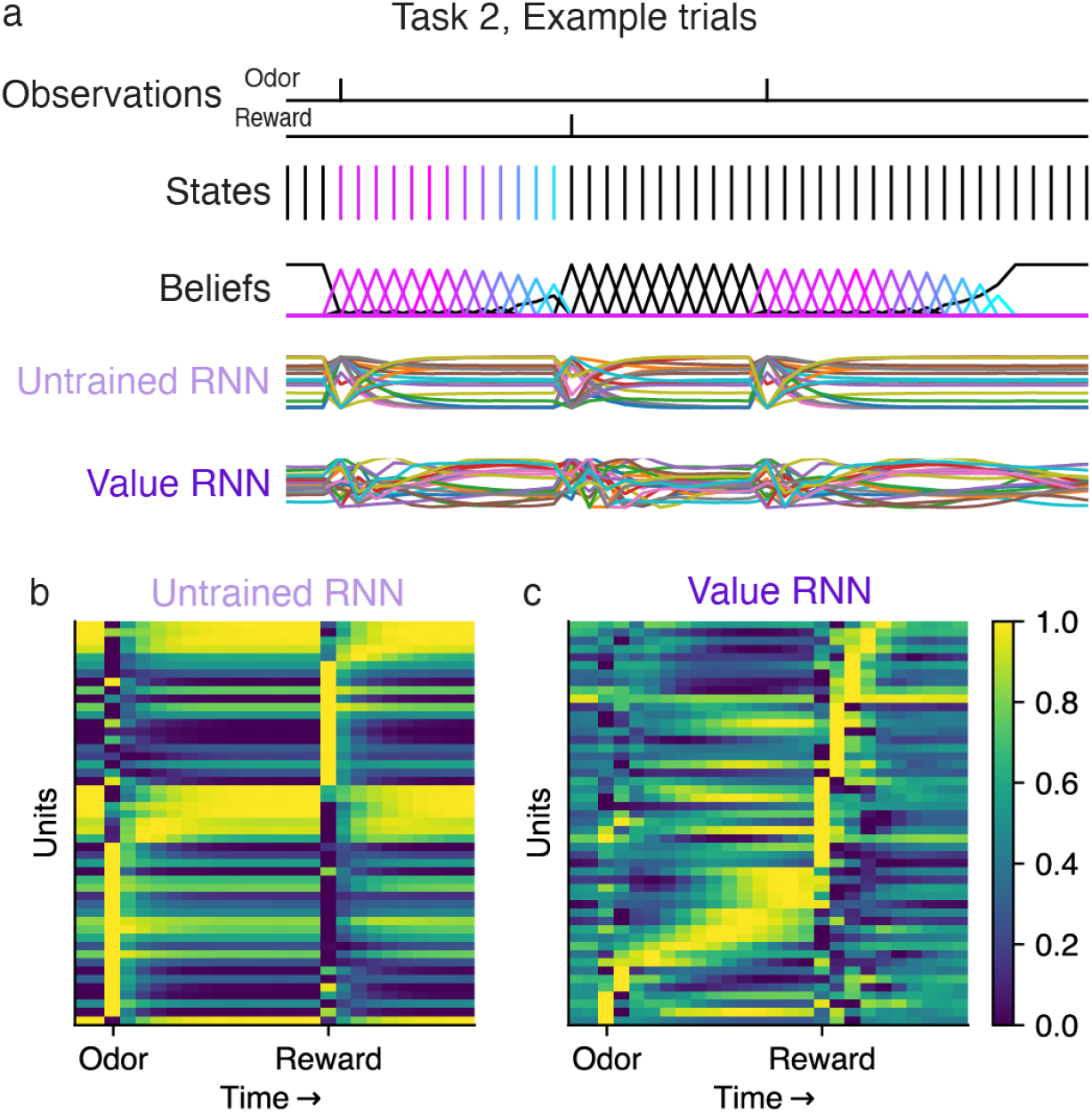
RNN activity before and after training on Starkweather Task 2. **a**. Observations, states, beliefs, and RNN activations on two example trials from Task 2. **b-c**. RNN unit activity (individually normalized to span between 0 and 1), with units sorted by time of peak activation on held-out trials, on an RNN before (panel **b**) and after (panel **c**) training. Both before and after after training, RNN units exhibited tuning to elapsed time following observations, with variance that scaled with elapsed time.

**Fig S2.**
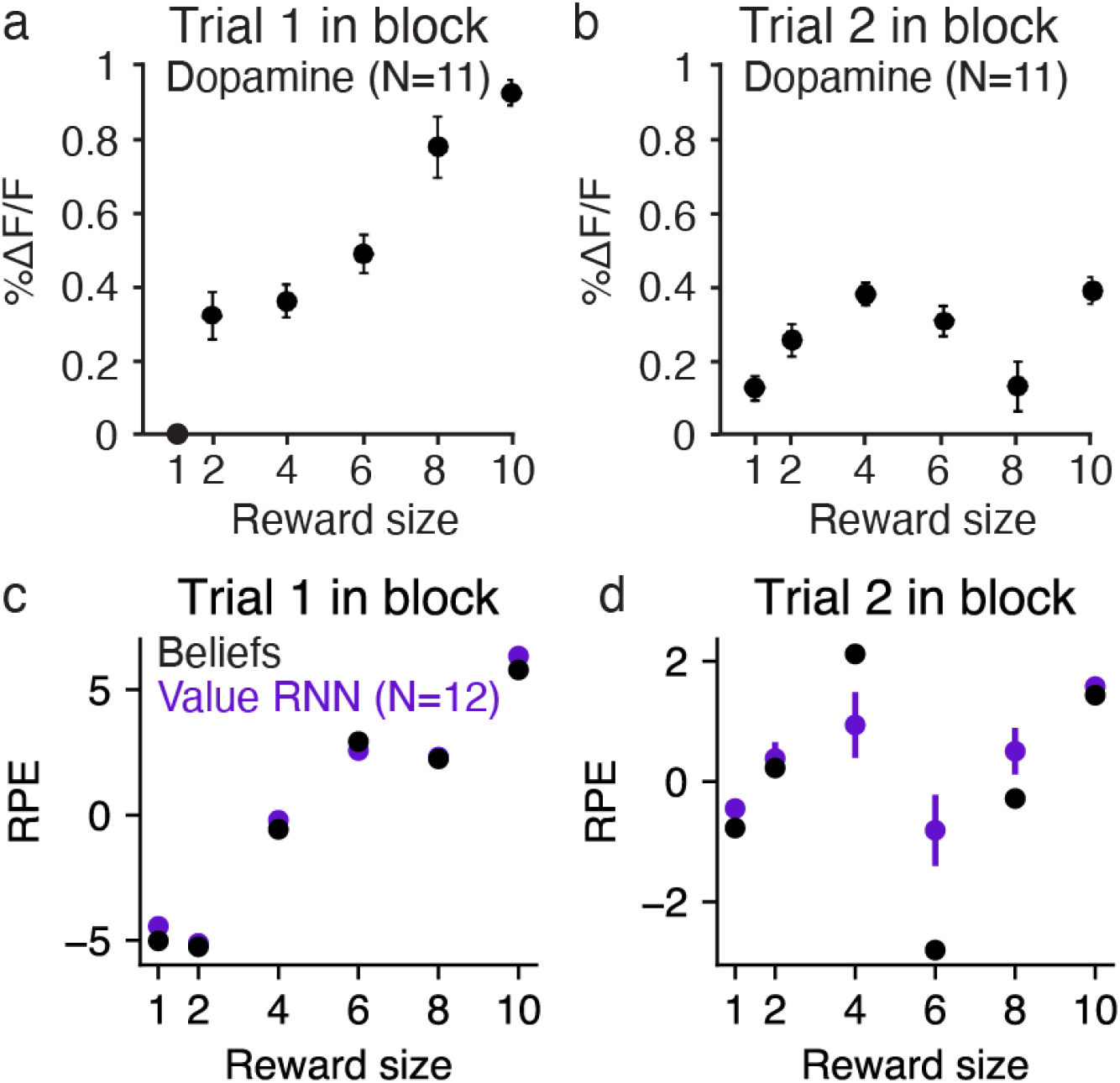
Value RNNs trained on the Babayan task recapitulate dopamine activity and belief RPEs in response to intermediate reward sizes. **a-b**. Average dopamine response on trial 1 (panel **a**) and trial 2 (panel **b**) during probe sessions including blocks with intermediate reward sizes. Circles and lines depict mean ±SE across *N* = 11 animals. Reproduced from Babayan et al. [10]. **c-d**. Same as panels **a-b**, but for the RPEs of the Belief model (black) and Value RNNs (purple). Value RNNs were trained on sessions including only blocks with rewards *r*_*t*_ ∈ {1, 10}, as in the main text. Value weights for the Belief model and Value RNNs were fit using a test session including 39 blocks each with *r*_*t*_ = 1 and *r*_*t*_ = 10, and 3 blocks each with *r*_*t*_ ∈ {2, 4, 6, 8}, similar to the proportions used in Babayan et al. [10]. RPEs were then measured on a different test session. Purple circles and lines depict mean ± SE across *N* = 12 models.

**Fig S3.**
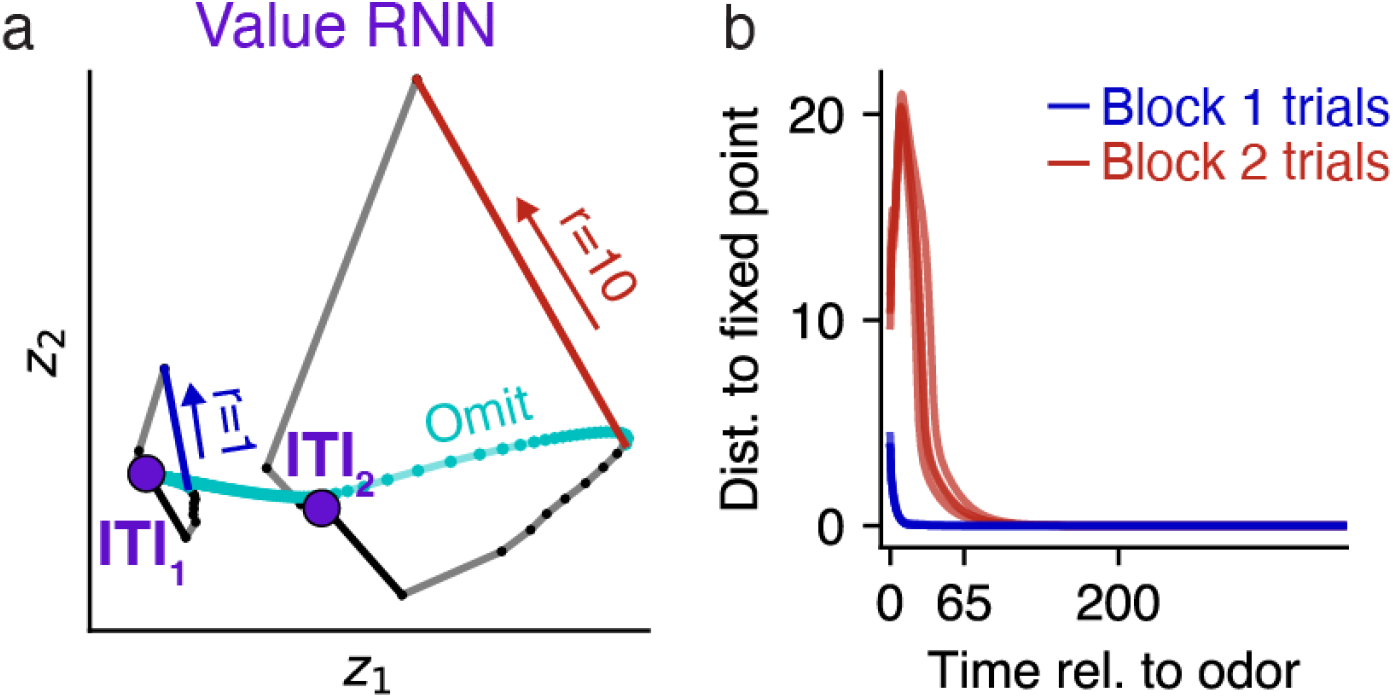
Value RNNs trained on the Babayan task exhibit one fixed point. **a**. RNN activity during two example trials, one during Block 1 (left) and the other during Block 2 (right). Same as Fig 7D. Here we also include RNN activity trajectories if each reward had been omitted. While activity for the Block 2 trial initially returns to the putative ITI_2_ state, it eventually returns to the true fixed point at ITI_1_ **b**. Distance of RNN activity from the single fixed point (e.g., ITI_1_ in panel **a**) following an odor observation (i.e., an omission trial). While the maximum ITI duration is theoretically infinite, the maximum ITI duration in the training data was at *t* = 65. RNN activity on Block 2 trials therefore remained separate from the activity on Block 1 trials for the range of experienced ITI durations.

## Notes

### Competing Interest Statement

The authors have declared no competing interest.

